# Subcellular interactions of neuropeptide Y and corticotropin-releasing factor in the central nucleus of the amygdala in the mouse

**DOI:** 10.1101/2025.09.19.677477

**Authors:** Himavarsha Yerraguntla, Joy Onyekachi, Laura L. Giacometti, Samuel L. Goldberg, Jessica R. Barson, Jacqueline M. Barker, Beverly A.S. Reyes

## Abstract

Neuropeptide Y (NPY) is ubiquitously distributed throughout the central nervous system. Recognized as a mediator of stress resilience, NPY has been shown to counteract the excitatory effects of the neuropeptide corticotropin-releasing factor (CRF), that orchestrates the stress response. In the mouse, while NPY and CRF exhibit a high degree of neuroanatomical association in the central nucleus of the amygdala (CeA) indicating potential significant interactions, the synaptic organizations of these neuropeptides have not been elucidated. In the present study, we determined the interactions between NPY and CRF in the CeA. Immunofluorescence microscopy presented that NPY-immunoreactive varicose processes were distributed throughout the CeA and contacted CRF-containing neurons. Using electron microscopy, immunoperoxidase labeling for NPY and gold-silver labeling for CRF showed that NPY-labeled axon terminals (NPY-t) form synapses with CRF-labeled dendrites (CRF-d). Semi-quantitative analysis revealed that 163 of NPY-t directly target CRF-d. In addition, approximately 85% of NPY-t form symmetric synapses with CRF-d while approximately 1% form asymmetric synapses. These findings provide the first ultrastructural evidence that NPY-containing axon terminals make direct contact with CRF-containing dendrites in the CeA. This suggests that the CRF-containing neurons in the CeA may be a key site for NPY action, potentially influencing brain regions involved in stress responses and stress-related psychiatric disorders, and alcohol use disorders.

## Introduction

Stress-related psychiatric disorders, including anxiety disorders, remain a pressing global health concern affecting the lives of millions each year (https://www.who.int/news-room/fact-sheets/detail/anxiety-disorders, accessed 17 August 2025). As such, the Global Burden of Diseases, Injuries, and Risk Factors Study (GBD) 2019 identifies anxiety disorders as among the most prevalent mental health conditions (GBD Collaborators, 2022) and the most commonly occurring category of psychiatric disorders in the United States (Kessler *et al*., 2005) and in some European countries (Alonso *et al*., 2007). In the United States alone, stress-related psychiatric disorders contribute to considerable economic losses due to their impact on individuals’ quality of life (Barbosa-Camacho *et al*., 2022). The central nucleus of the amygdala (CeA) is one of the key brain nuclei implicated in stress-related psychiatric disorders and has emerged as a critical target for developing novel psychopharmacological therapies given its central role in emotional processing, arousal and the stress response (Adhikari, 2014; Paretkar & Dimitrov, 2018). The CeA functions as the primary output station of the amygdala and is extensively connected to the endocrine and autonomic centers in the hypothalamus and brainstem, and thereby mediates both behavioral and autonomic responses to emotionally arousing stimuli (Tsukioka *et al*., 2022). Dysregulation of the CeA has been observed in anxiety related to stress, substance use and withdrawal (Patel *et al*., 2022).

Corticotropin-releasing factor (CRF) is a 41-amino acid neuropeptide first identified by Vale and colleagues in 1981 (Vale *et al*., 1981). It plays a prominent role in the orchestration of endocrine, autonomic and behavioral responses to stress. Dysregulation of CRF is a significant component and a key factor in the development of stress-related psychiatric disorders. Importantly, overactivation of the CRF system is an essential feature of the physiological and emotional responses to stress (Tovote *et al*., 2015). While CRF-containing neurons are densely distributed in the hypothalamus, as first reported (Vale et al., 1981), CRF-synthesizing cell bodies are also abundantly distributed in the CeA, which is considered a major source of extra-hypothalamic CRF in the brain. Experimental knockdown of CRF in the CeA significantly diminishes the stress-induced anxiety (Ventura-Silva et al., 2020).

Similar to CRF, neuropeptide Y (NPY) is also enriched in the CeA. NPY is a 36-amino acid peptide that acts as a neuromodulator of the stress response but, in contrast to CRF, it is associated with resiliency to adverse effects of stress (Sah et al., 2014) and is considered an anti-stress neuropeptide. In fact, Sah and colleagues termed NPY as the resilience-to-stress factor in humans (Sah & Geracioti, 2013; Sah *et al*., 2014). Anatomical investigations have shown a dense plexus of NPY-immunoreactive processes in the CeA (Gustafson *et al*., 1986).

Previous anatomical studies have demonstrated the localization of the NPY and CRF in the CeA (Gustafson *et al*., 1986; Reyes *et al*., 2017; Reyes *et al*., 2019; Van Bockstaele *et al*., 1998; Van Bockstaele *et al*., 1999) while functional studies, such as those on alcohol use disorder (AUD) (Gilpin et al., 2015), demonstrate an association between NPY and CRF. These studies suggest interactions between NPY and CRF in the CeA; however, it remains to be investigated whether NPY and CRF form a direct cellular substrate of interactions. High-resolution imaging such as electron microscopy is essential to elucidate this relationship at the ultrastructural level. Therefore, in the current study, we employed immunohistochemistry, immunofluorescence and immunoelectron microscopy to identify anatomical distribution of NPY and CRF in the CeA. We investigated the interaction between NPY and CRF and observed that NPY innervates CRF-containing neurons in the CeA, providing the structural evidence for a potential synaptic interface underlying their functional interplay in stress-related psychiatric disorders and AUD.

## Materials and methods

The procedures employed in the present study were approved by the Institutional Animal Care and Use Committee of Drexel University and adhered to the National Institute of Health’s Guide for the Care and Use of Laboratory Animals. Every effort was made to use the minimum number of animals required to obtain scientifically reliable data, and all measures were taken to minimize any animal distress.

### Experimental animals

Eight adult male C57BI/6J mice (Jackson Laboratory, Sacramento, CA) were used in this study. They were housed five mice per cage on a 12-hour light/dark cycle (lights on at 7:00 am) in a temperature-controlled (25°C) colony room. They were given *ad libitum* access to food (LabDiet 5053; PicoLab Rodent Diet, St. Louis, MO) and water. These mice were used for examining the cellular association of NPY and CRF-containing profiles in the CeA.

### Immunofluorescence

The region analyzed included primarily the CeA that exhibits abundance in CRF immunoreactivity as previously described (Asan et al., 2005; Wang et al., 2011). At the level of bregma -0.58 to -1.82 mm, the CeA, spanning from the cranial to caudal end, is bounded dorsally by the interstitial nucleus of the posterior limb of the anterior commissure, basal nucleus of Meynert and lateral globus pallidus, laterally by the intercalated nuclei of the amygdala, ventral endopiriform nucleus and basolateral amygdaloid nucleus (anterior), medially by the basal nucleus of Meynert, substantia innominata, medial amygdalar nucleus, (anterodorsal) and ventrally by the basomedial amygdaloid nucleus (anterior part), anterior amygdaloid area (dorsal), intercalated amygdaloid nucleus (main) and optic chiasm (Franklin & Paxinos, 2007).

Four C57BL/6J adult male mice were used for immunofluorescence portion of the study. One of the four mice was naive, while the remaining three received bilateral microinjections (0.2 μl per side, 0.1µl/min, 5 min diffusion) of pAAVrg-hSyn-GFP (Addgene, Watertown, MA) in the locus coeruleus (anterior-posterior to lambda -0.9 mm; medial-lateral ± 0.9 mm; dorsal-ventral -2.9 mm). To minimize the use of additional animals, these three mice were sourced from the control group of a separate experiment, the results of which are not included in this manuscript. Following recovery for 1-2 weeks, the three mice then underwent forced air exposure for four cycles. One cycle consisted of 16 h/day of forced air inhalation for 4 days and each cycle was separated by 72 h. Mice received an intraperitoneal injection of 1 mmol pyrazole in saline prior to being placed in the air chamber. The mouse brains were used exclusively to assess whether a cellular interaction exists between NPY and CRF expression in the CeA, utilizing confocal microscopy. All four mice examined through confocal imaging showed no observable variation in cellular interaction between NPY and CRF expression.

These four mice were deeply anesthetized with isoflurane (Abbott Laboratories, Abbott Park, IL). They were transcardially perfused through the ascending aorta with heparin followed by 4% formaldehyde in 0.1 M phosphate buffer (PB; pH 7.4) using the Masterflex peristaltic pump (Cole-Parmer, Vernon Hills, IL). The brains were removed, blocked, cut in half in coronal orientation, immersed in 4% formaldehyde overnight at 4°C, and stored in 30% sucrose solution in 0.1 M PB at 4°C for two to three days. A sequential set of 40 µm coronal sections through the rostrocaudal extent of each brain was collected using a cryostat (Microm HM 50, Microm International GmbH, Walldorf, Germany). Serial sections through the CeA were incubated for 30 min in 1% sodium borohydride in 0.1 M PB to reduce amine-aldehyde complexes thereby diminishing autofluorescence (Clancy and Cauller, 1998). The tissue sections were then incubated in 0.5% bovine serum albumin (BSA) in 0.1 M tris-buffered saline (TBS; pH 7.6) for 30 min. Following extensive rinsing in 0.1 M TBS, the tissue sections were incubated in a cocktail containing rabbit anti-NPY at 1:4,000 (Sigma-Aldrich, Inc., St. Louis, MO) and guinea-pig anti-CRF at 1:2,000 (Peninsula Laboratories, San Carlos, CA). Incubation time was 15-18 h on a rotary shaker at room temperature. Sections were then washed in 0.1 M TBS and incubated in a secondary antibody cocktail containing fluorescein isothiocyanate (FITC) donkey anti-rabbit (1:200; Jackson ImmunoResearch Laboratories Inc., West Grove, PA) and tetramethyl rhodamine isothiocyanate (TRITC) donkey anti-guinea pig (1:200; Jackson ImmunoResearch) antibodies prepared in 0.1 % BSA and 0.25% Triton X-100 in 0.1 M TBS for 2 h in the dark on a rotary shaker. Control tissues were processed in parallel, one with the omission of primary antibodies and the other with the omission of secondary antibodies. The tissue sections were visualized using an Olympus 1x81 inverted microscope (Olympus, Hatagaya, Shibuya-Ku, Tokyo, Japan) equipped with lasers (Helium Neon laser and Argon laser; models GLG 7000; GLS 5414A and GLG 3135, Showa Optronics Co., Tokyo, Japan) excitation wavelength of 488 and 543. The microscope is also equipped with filters (DM 405-44; BA 505-605; and BA 560-660) with Olympus Fluoview ASW FV1000 program (Olympus, Hatagaya, Shibuya-Ku, Tokyo, Japan). Figures were assembled and adjusted for brightness and contrast in Adobe Photoshop.

### Immunoelectron microscopy

Four C57BL/6J adult male mice were deeply anesthetized with isoflurane (Abbott Laboratories) and transcardially perfused through the ascending aorta with heparin, followed with 3.75% acrolein (Chem Service, Inc., West Chester, PA) and 2% formaldehyde in 0.1 M PB. The brains were removed, cut in half in coronal orientation, and immersed in 2% formaldehyde overnight at 4°C. Forty-μm-thick coronal sections through the rostrocaudal extent of the CeA were cut on a vibratome (Pelco EasiSlicer, Ted Pella Inc., Redding, CA). Tissue sections were processed for electron microscopy as previously described (Reyes, et al., 2011; Reyes et al., 2017; Reyes et al., 2019). Briefly, tissue sections were placed for 30 min in 1% sodium borohydride in 0.1 M PB to remove reactive aldehydes and incubated in 0.5% BSA in 0.1 M TBS for 30 min. Following thorough rinses in 0.1 M TBS, tissue sections were incubated overnight in a cocktail containing rabbit anti-NPY at 1:2,000 (Sigma-Aldrich, Inc.) and guinea-pig anti-CRF at 1:1,000 (Peninsula Laboratories) with no Triton X-100 in the solution. The following day, tissue sections were rinsed three times in 0.1 M TBS and incubated in biotinylated donkey anti-rabbit (1:400; Vector Laboratories, Burlingame, CA) for 30 min followed by rinses in 0.1 M TBS. Subsequently, a 30-minute incubation of avidin-biotin complex (Vector Laboratories) followed. For all incubations and washes, sections were continuously agitated with a rotary shaker. NPY was visualized by a 5-minute reaction in 22 mg of 3,3’-diaminobenzidine (Sigma-Aldrich Inc.) and 10 µl of 30% hydrogen peroxide in 100 ml of 0.1 M TBS. Immunoperoxidase labeling was used to visualize NPY immunoreactivity.

While NPY immunoreactivity was labeled with immunoperoxidase, CRF immunoreactivity was identified with immunogold-silver labeling. For gold-silver localization of CRF, sections were rinsed three times with 0.1 M TBS, followed by rinses with 0.1 M PB and 0.01 phosphate buffered saline (PBS; pH 7.4). Sections were then incubated in a 0.2% gelatin-PBS and 0.8% BSA buffer for 10 min. This was followed by incubation in goat anti-rabbit IgG conjugate in 1 nm gold particles (1:50; Electron Microscopy Sciences, Hatfield, PA) at room temperature for 2 h. Sections were then rinsed in buffer containing the same concentration of gelatin and BSA as above and subsequently rinsed with 0.01 M PBS. Then sections were incubated in 2% glutaraldehyde (Electron Microscopy Sciences) in 0.01 M PBS for 10 min followed by washes in 0.01 M PBS and 0.2 M sodium citrate buffer (pH 7.4). A silver enhancement kit (Aurion R-GENT SE-EM kit, Electron Microscopy Sciences) was used for silver intensification of the gold particles. Following intensification, tissue sections were rinsed in 0.2 M citrate buffer and 0.1 M PB and incubated in 2% osmium tetroxide (Electron Microscopy Sciences) in 0.1 M PB for 1 h, washed in 0.1 M PB, dehydrated in an ascending series of ethanol followed by propylene oxide and flat embedded in Epon 812 (Electron Microscopy Sciences). Thin sections were cut at 70 nm in thickness with a diamond knife (Diatome-US, Fort Washington, PA) using a Jeol Ultracut E (Jeol Ltd., Peabody, MA). Sections were examined with an electron microscope (Tecnai G2, Fei Company, Hillsboro, OR), with digital images captured by an AMT Base Model: BIOSPR12 camera system (Advance Microscopy Techniques Corp., Danvers, MA). Figures were assembled and adjusted for brightness and contrast in Adobe Photoshop.

### Specificity of antisera

The NPY antibody was raised in rabbit. The specificity of the NPY antibody in the present study was previously reported using brain sections from mice with a deletion of NPY (NPY-KO) and wildtype (WT) control (Theisen *et al*., 2018). NPY immunoreactivity was only observed in WT control while the absence of NPY immunoreactivity was evident in NPY-KO mice (Theisen *et al*., 2018). The CRF antibody was raised in guinea pig and was generated against human/mouse/rat CRF peptide (SEEPPISLDLTFHLLREVLEMARAEQLAQQAHSNRKLMEII) (Peninsula Laboratories). The specificity of CRF immunolabeling was examined and tested by absence of immunoreactivity in areas not known to express CRF as we previously reported (Reyes et al., 2017). Moreover, it was previously reported by Sawada and colleagues that preabsorption with the antigenic CRF peptide eliminated CRF immunoreactivity (Sawada *et al*., 2008). Lastly, sections processed in the absence of the primary CRF antiserum did not exhibit CRF immunoreactivity (Reyes *et al*., 2017; Theisen *et al*., 2018; Reyes *et al*., 2019). While tissue sections were incubated with the primary antibodies, NPY and CRF, some sections were processed in parallel and served as negative control groups that do not contain primary antibody. Sections processed in the absence of NPY or CRF primary antibody did not exhibit any detectable immunoreactivity. Some sections were also processed with primary antibody but no secondary antibody and used to reveal any endogenous reactivity to primary antibodies. To evaluate cross-reactivity of the primary antisera by secondary antisera, some sections were processed for dual labeling with omission of one of the primary antisera. Tissue sections for electron microscopy were obtained from mice with optimal preservation of ultrastructural morphology and with both markers clearly apparent were used for the analysis.

### Electron microscopy data analysis

We have previously described the semi-quantitative approach used in the present study (Bangasser *et al*., 2013; Reyes *et al*., 2005; Reyes *et al*., 2011; Reyes *et al*., 2017). Although acrolein fixation is known to enhance the preservation of ultrastructural morphology, it also presents certain limitations. These limitations include restricted or uneven penetration of immunoreagents in thick tissue sections (Chan *et al*., 1990). Consequently, the restricted penetration of NPY and CRF may result in an underestimation of the relative distribution frequencies. To overcome this limitation, we restricted our sampling to the tissue sections adjacent to the tissue-Epon interface where reagent penetration is most effective. Additionally, we found that limiting section collection to the surface layer reduces potential artifacts linked to insufficient antisera penetration. The analysis of tissue sections collected at the plastic-tissue interface ensured that both markers were detectable in all sections used for analysis (Bangasser *et al*., 2013; Reyes *et al*., 2005; Reyes *et al*., 2011; Reyes *et al*., 2017). At least six grids containing ultrathin sections were collected from individual nonadjacent 40 μm-thick amygdalar Vibratome sections. At least two Vibratome sections were examined on each animal. All axon and axon terminals labeled with immunoperoxidase for NPY were photographed and classified when immunogold-silver labeling for CRF was also present in the neuropil, terminals or dendrites within the same fields, at a minimum magnification of 6,500X. The cellular elements were identified based on the description of Peters and colleagues (Peters & Palay, 1996). Somata contained a nucleus, Golgi apparatus and smooth endoplasmic reticulum. Proximal dendrites contained endoplasmic reticulum, were postsynaptic to axon terminals and were larger than 0.7 μm in diameter. A terminal was considered to form a synapse if it showed a junctional complex, a restricted zone of parallel membranes with slight enlargement of the intercellular space, and/or associated with postsynaptic thickening. Asymmetric synapses were identified by thick postsynaptic densities (Gray’s type I; Gray, 1959); in contrast, symmetric synapses had thin densities (Gray’s type II; Gray, 1959) both pre- and postsynaptically. Undefined contacts were characterized by a junctional complex that was not readily identifiable. A non-synaptic contact or apposition was defined as an axon terminal plasma membrane juxtaposed to that of a dendrite or soma devoid of recognizable membrane specializations and no intervening glial processes.

### Identification of immunogold-silver labeling

CRF was visualized using immunogold-silver enhancement. Selective gold-silver labeled profiles were identified by the presence, in single thin sections, of at least two gold particles within a cellular compartment. Profiles containing CRF-labeled axon terminals were counted and their association with NPY was determined. As we previously reported, a single spurious immunogold-silver labeling can contribute to false-positive labeling and can be detected in blood vessels, myelin or nuclei (Reyes *et al*., 2005; Reyes *et al*., 2011; Reyes *et al*., 2017). Minimal spurious labeling was encountered in the present study, therefore the criterion for defining a process exhibiting immunogold-silver labeling was set by the presence of at least two silver particles in a profile. A profile containing only one gold particle in adjacent thin sections was designated as lacking detectable immunoreactivity.

## Results

### Immunofluorescence labeling of CRF and NPY

Immunofluorescence labeling using FITC and TRITC-tagged secondary antibodies to identify CRF and NPY was conducted at approximately the same AP coordinates through the CeA (Fig. 1A-C). While CRF and NPY immunoreactivities in the CeA have been previously described in separate neuroanatomical studies in mice (Asan et al., 2005; Wang et al., 2011; Wood et al., 2016), the existence of a cellular substrate of interactions between these neuropeptides has not been investigated. Our present results demonstrate that CRF immunoreactivity is localized in neurons and fibers within the CeA (Fig. 1A, C), consistent with the previous reports in mice using immunohistochemistry (Asan et al., 2005; Wang et al., 2011). Similarly, NPY immunoreactive neurons and abundant NPY-immunoreactive fibers were observed in the CeA (Fig. 1B, C), in agreement with prior findings in mice using immunofluorescence (Wood et al., 2016). NPY immunoreactive processes (red labeling) were observed contacting CRF-labeled neurons (green labeling) in the CeA (Fig. 1A-C). The interaction between NPY and CRF suggests that some NPY-immunoreactive processes may innervate and impact the activity of CRF-expressing neurons. However, not all NPY-immunoreactive processes formed contacts with CRF-labeled neurons (Fig. 1A-C).

**Figure 1.**
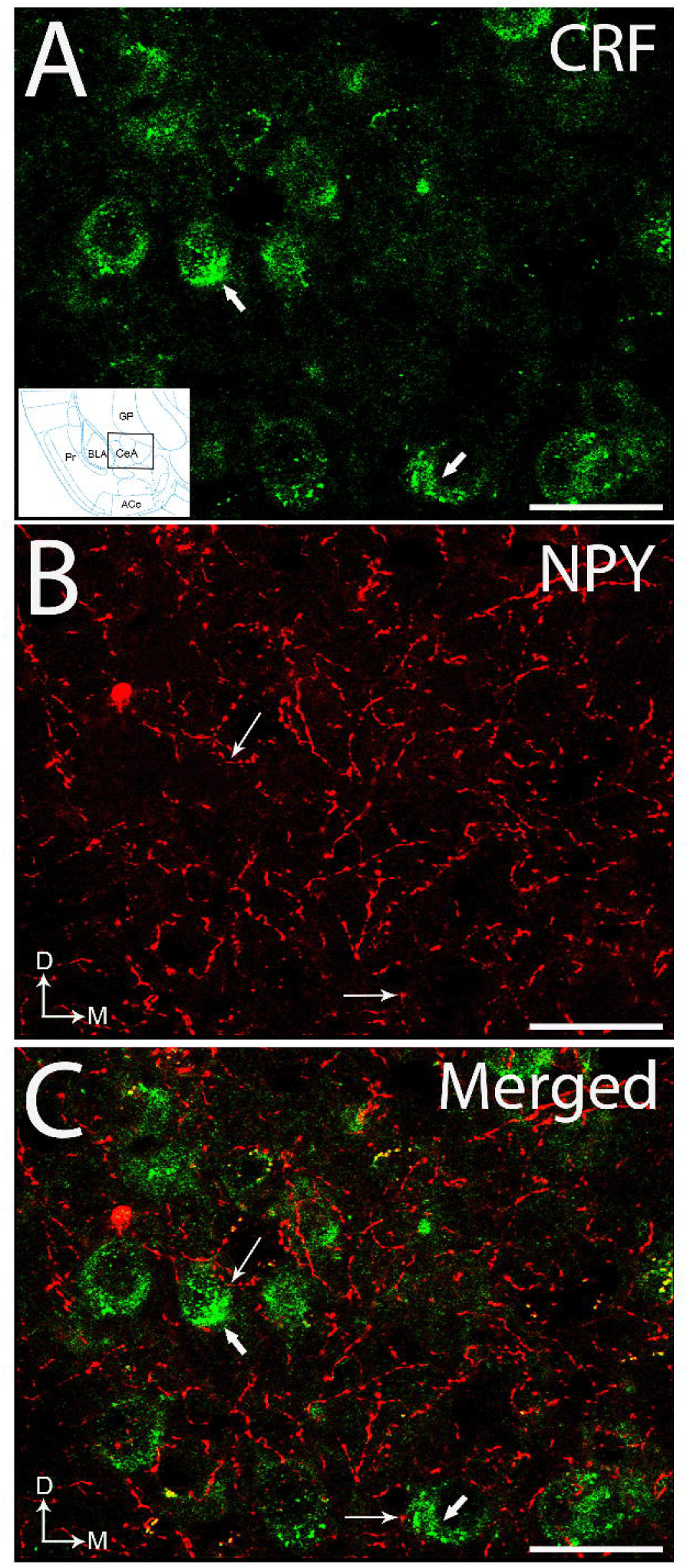
Confocal fluorescence photomicrographs showing immunolabeling of neuropeptide Y (NPY) with respect to corticotropin-releasing factor (CRF) in the central nucleus of the amygdala (CeA). NPY-containing processes are contacting CRF-labeled neurons in the CeA. **A.** Fluorescence photomicrograph showing CRF immunolabeling detected using a fluorescein isothiocyanate-conjugated secondary antibody (FITC donkey anti-guinea pig; green). Thick arrows point to CRF-labeled neurons. Inset shows a schematic diagram adapted from the mouse brain in stereotaxic coordinates (Franklin & Paxinos, 2007) showing the anterior posterior (anterior-posterior to bregma -0.94 mm) level of the region of the central nucleus of the amygdala where the photomicrograph was obtained. **B.** Fluorescence photomicrograph showing NPY immunolabeling detected using a rhodamine isothiocyanate-conjugated secondary antibody (TRITC donkey anti-rabbit; red) can be identified in the processes. Thin arrows indicate individual NPY-labeled processes. Arrows indicate dorsal (D) and medial (M) orientation of the sections. **C.** A merged image of A and B showing NPY-labeled processes contacting CRF-labeled neurons in the CeA. Thick arrows point to CRF-labeled neurons while thin arrows indicate individual NPY-labeled processes. Arrows indicate dorsal (D) and medial (M) orientation of the sections. Aco, anterior optical amygdaloid nucleus; BLA, basolateral nucleus of the amygdala; CeA, central nucleus of the amygdala; GP, globus pallidus; Pr, piriform cortex; Scale bar, 100 μm.

### Ultrastructural analysis of CRF and NPY interactions in the CeA

Given that the dual immunofluorescence demonstrated that NPY-immunoreactive processes are contacting CRF-containing neurons, we proceeded to use electron microscopy to closely examine and quantify the cellular substrate of interactions between the NPY and CRF. At the ultrastructural level, immunoperoxidase labeling for NPY and immunogold-silver labeling for CRF were localized in the same section within the neuropil of the CeA sampled for the semi-quantitative analysis (Fig. 2, 4). Electron photomicrographs showed that NPY and CRF immunoreactivities were richly distributed throughout the neuropil (Fig. 2, 4). The immunoperoxidase labeling indicative of NPY demonstrated an electron diffuse reaction product within vesicles filled axon terminals, and distinguishable from the immunogold-silver deposits indicative of CRF (Fig. 2, 4). Silver intensified gold immunolabeling for CRF appeared as irregularly shaped black deposits that were distributed within the cytoplasm and along the plasma membrane (Fig. 2, 4). The region sampled in the CeA for immunoelectron microscopy was similar to that demonstrated for immunofluorescence microscopy (Fig. 1). Both NPY and CRF immunoreactivities were abundant in the neuropil at the ultrastructural level (Fig. 2, 4). The organelles within the axon terminals were sometimes obscured by the dense immunoperoxidase product for NPY immunolabeling (Fig. 2, 4). However, in axon terminals that were lightly labeled, NPY-labeled axon terminals contained distinct vesicular populations characterized by small clear spherical vesicles (Fig. 2, 4). The majority of NPY-immunoreactive profiles was identified primarily in unmyelinated axon terminals (Fig. 2, 4) while CRF immunoreactivity was identified in dendritic processes (Fig. 2, 4-5), soma (Fig. 6C), and a few cases in axon terminals (Fig. 6A). Thus, the findings of dual immunofluorescence in the CeA (Fig. 1) are confirmed at the ultrastructural level (Fig. 2, 4-6). Presented in Fig. 2, 4-5 are NPY axon terminals contacting CRF-labeled dendrites. Single CRF-containing dendrites in the CeA received convergent input from NPY-labeled axon terminals (Fig. 4C). Convergent input was characterized by synaptic contacts formed by multiple NPY-labeled axon terminals synapsing on a common CRF-labeled dendrite on the same cellular profile (Fig. 4C). Semiquantitative analysis revealed that of the 334 axon terminals demonstrating NPY immunoreactivity in the neuropil, 49% (n = 163) directly contacted dendrites exhibiting CRF-immunoreactivity. When synaptic specializations were identifiable between NPY-immunoreactive axon terminals and dendrites containing CRF, they were morphologically heterogeneous (Fig. 2, 4-5). When synaptic specialization was recognizable they were either symmetric, indicative of inhibitory synapses (Gray’s type II; (Gray, 1959), or asymmetric, representating excitatory synapses (Gray, 1959). However, some NPY-labeled axon terminals that contacted CRF-containing dendrites could not be clearly classified as either type and were therefore categorized as undefined. Of the 163 NPY-axon terminals in direct contact with CRF-containing dendrites that formed synaptic contacts, 85% (138/163; Fig. 3A) were symmetric type (Fig. 3A, 4A-B) while 1% (2/163; Fig. 3B) formed asymmetric synapses. NPY-labeled axon terminals that formed symmetric synapses were characterized by thin postsynaptic densities, while asymmetric synapses were characterized by thick postsynaptic densities. Some NPY-labeled axon terminals lacked clearly distinguishable synaptic specializations with post-synaptic targets (Fig. 3C, 4-B, D). About 14% (23/163; Fig. 3C) of NPY-labeled axon terminals did not form recognizable synapses in the planes of section analyzed (Fig. 3C, 4-B, D). Occasionally, NPY-labeled axon terminals were observed contacting a CRF-labeled soma (Fig. 5A). Some NPY-labeled axon terminals were co-localized with CRF (Fig. 5B). Approximately, 28% of NPY-axon terminals were co-localized with CRF (92/334; Fig. 5B). It was also observed that some NPY and CRF-immunoreactivities were found in a common dendrite (Fig. 5B-C). We also processed the reverse immunolabeling, where NPY was labeled with immunogold-silver particles while CRF was labeled with immunoperoxidase labeling. Figure 6 shows that NPY-labeled axon terminals form direct synaptic contact with CRF-labeled dendrites.

**Figure 2.**
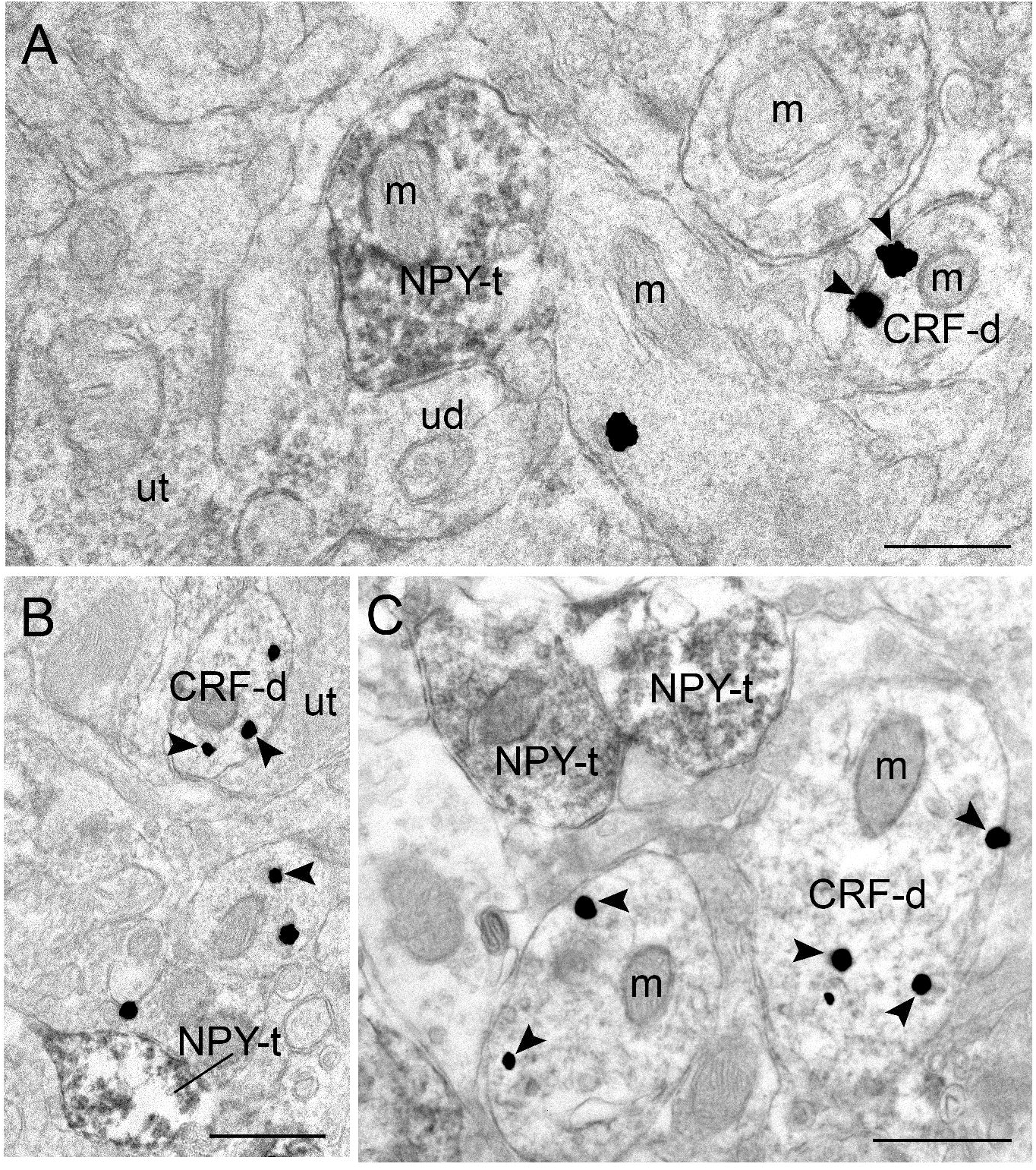
Electron photomicrographs showing immunoperoxidase labeling for neuropeptide Y (NPY) and immunogold-silver labeling for corticotropin-releasing factor (CRF) in the central nucleus of the amygdala (CeA) **A-B.** Sections showing an NPY-labeled axon terminal (NPY-t) contacting an unlabeled dendrite (ud). NPY immunolabeling is densely distributed within the axon terminal (NPY-t). CRF-containing dendrite (CRF-d) is labeled with immunogold-silver particles (arrows). **C.** Two NPY-labeled axon terminals (NPY-t) are adjacent to each other showing an immunoperoxidase NPY labeling that contains vesicles. In all panels A-C, NPY-t is shown in a separate profile from a CRF-d. Arrowheads point to immunogold-silver particles indicative of CRF immunoreactivity. Scale bars = A. 0.4 µm; B. 0.5 µm and C. 0.6 µm.

**Figure 3.**
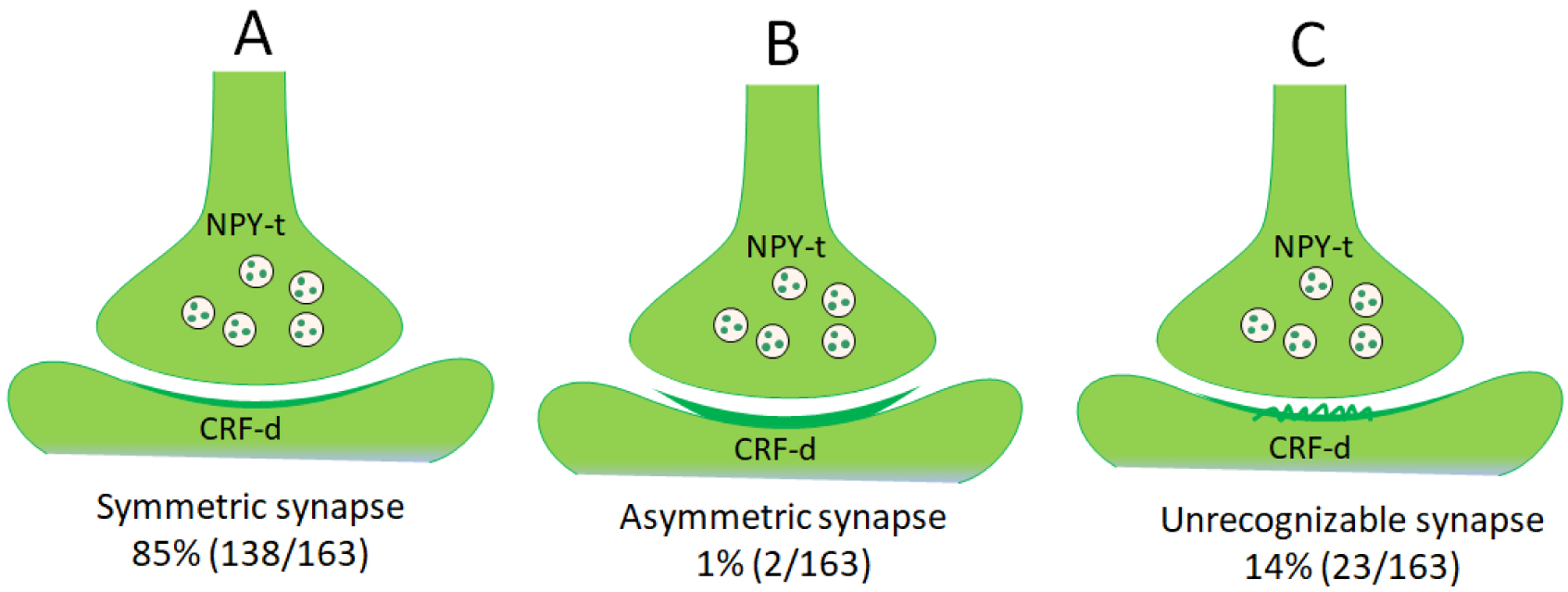
Schematics and quantifications of the cellular substrates of interactions between neuropeptide Y (NPY) and corticotropin-releasing factor (CRF) in the central nucleus of the amygdala. **A.** Symmetric synapse formed by NPY-containing axon terminals with CRF-containing dendrites. **B.** Asymmetric synapse formed by NPY-containing axon terminals with CRF-containing dendrites. **C.** Unrecognizable synaptic contacts formed by NPY-containing axon terminals with CRF-containing dendrites.

**Figure 4.**
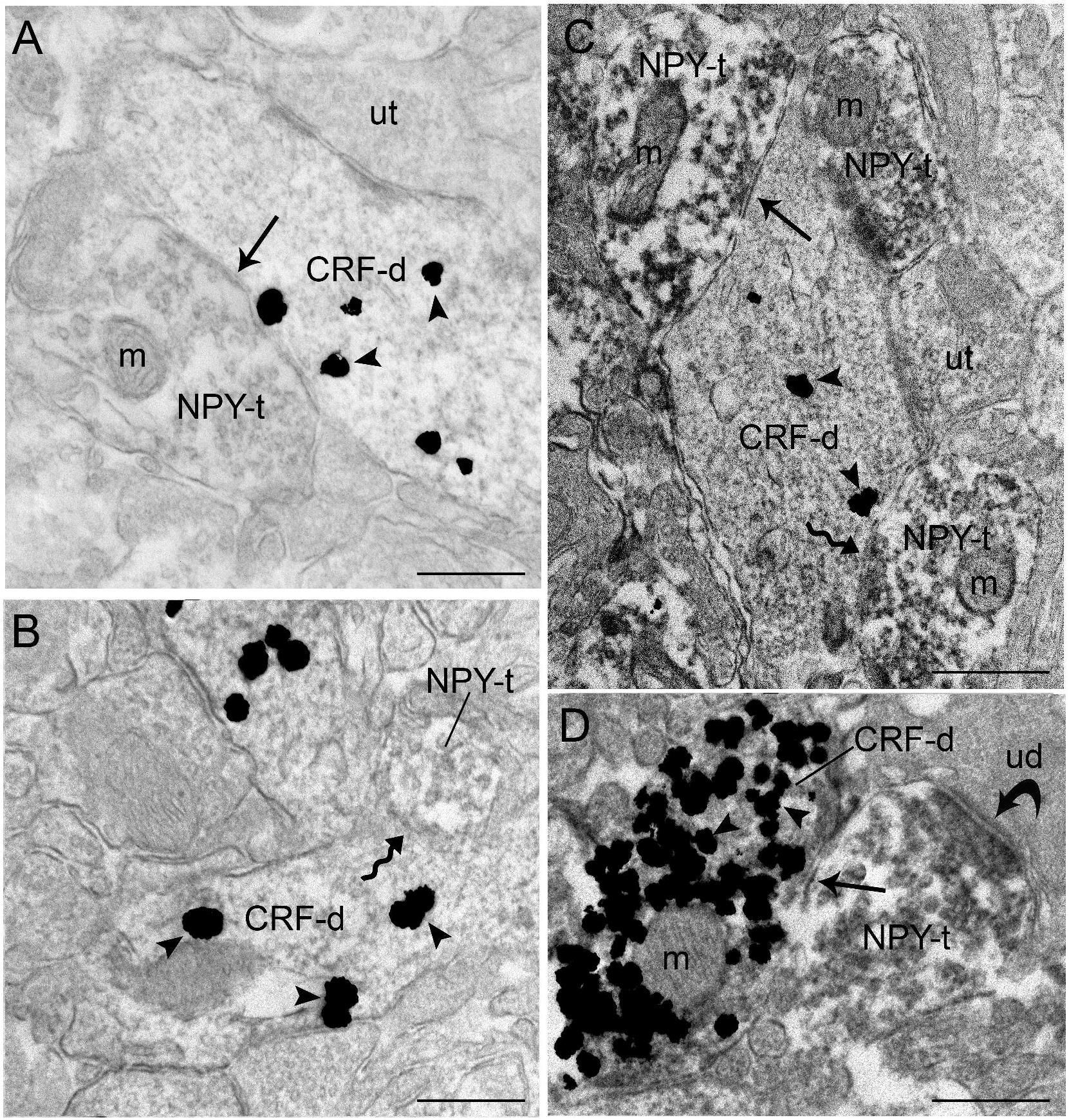
Electron photomicrographs showing neuropeptide Y (NPY) forming synaptic specializations with dendrites containing CRF in the central nucleus of the amygdala (CeA) **A.** NPY-labeled axon terminal (NPY-t) forms symmetric type synaptic contact (arrow) with CRF-labeled dendrite (CRF-d). The NPY-labeled axon terminal contains numerous small clear vesicles. Scale bar = 0.4 µm. **B.** NPY-t forms an unrecognizable synaptic contact with a CRF-d (curved arrow). Scale bar = 0.4 µm. **C.** Multiple NPY-t converge onto a single CRF-d. NPY-t on top forms a symmetric synapse (arrow) while the NPY-t below forms unrecognizable synaptic contact. Scale bar = 0.6 µm. **D.** NPY-t forms a symmetric contact (arrow) with a CRF-d and a symmetric contact (curved arrow) with unlabeled dendrite (ud). Arrowheads point to immunogold-silver particles indicative of CRF immunoreactivity. ut, unlabeled axon terminal; m, mitochondria.

**Figure 5.**
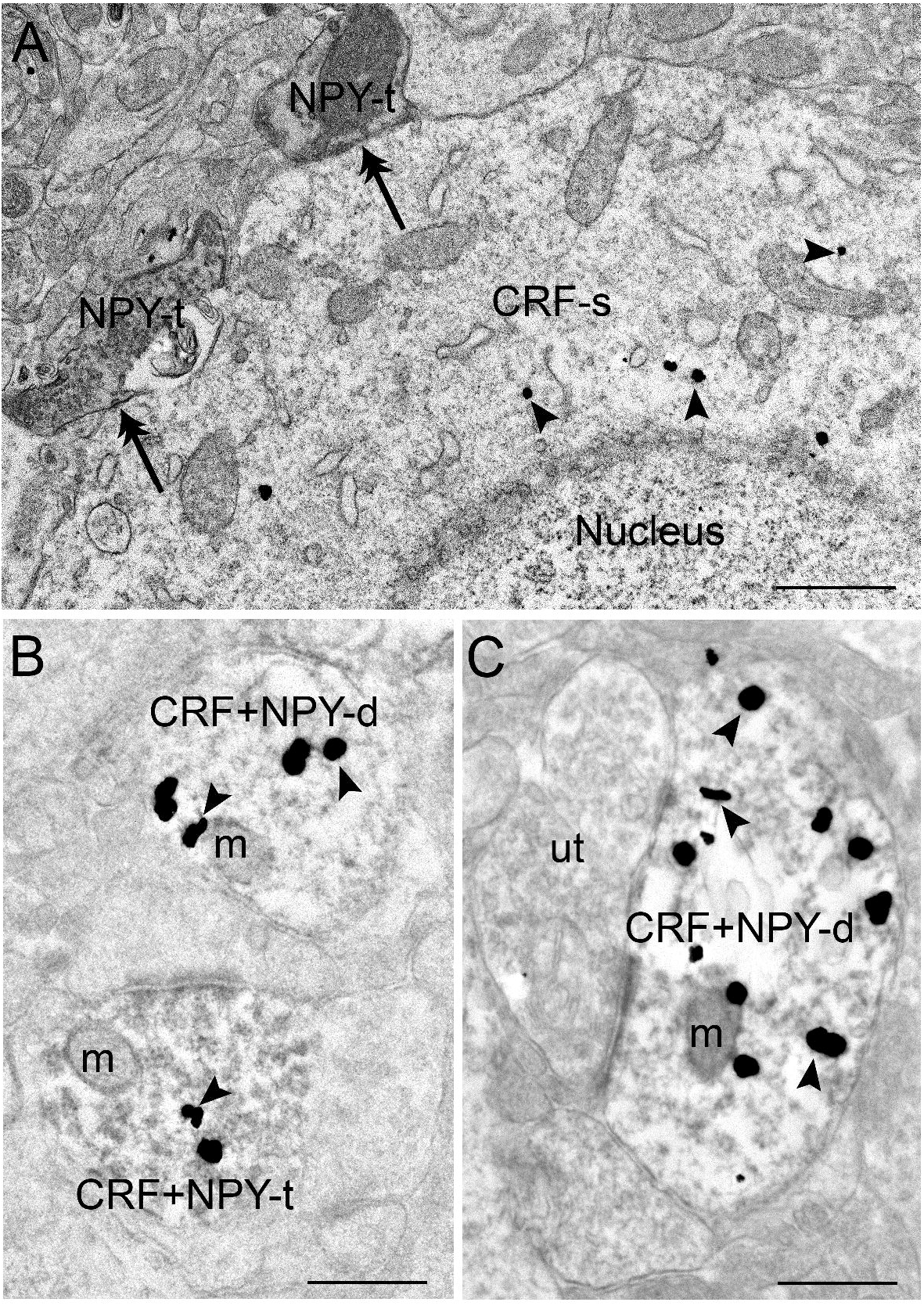
**A**. Electron photomicrograph showing neuropeptide Y (NPY) processes that form synaptic contact with a perikaryon containing CRF in the central nucleus of the amygdala (CeA). NPY-labeled axon terminals (NPY-t) form synaptic contact (double arrows) with CRF-labeled soma (CRF-s). Arrows point to immunogold-silver particles indicative of CRF immunoreactivity. Scale bar = 0.6 µm. **B.** Electron photomicrograph showing co-localization of neuropeptide Y (NPY) and CRF in a common axon terminal that forms synaptic specializations with an unlabeled dendrite (ud) in the central nucleus of the amygdala (CeA). The CRF and NPY are co-localized in a dendrite on top (CRF+NPY-d). Arrows point to immunogold-silver particles indicative of CRF immunoreactivity. Scale bar = 0.4 µm. **C.** Unlabeled axon terminal (ut) contacts a dendrite showing NPY and CRF immunoreactivities (CRF+NPY-d). Arrowheads point to immunogold-silver particles indicative of CRF immunoreactivity Scale bar = 0.4 µm

**Figure 6.**
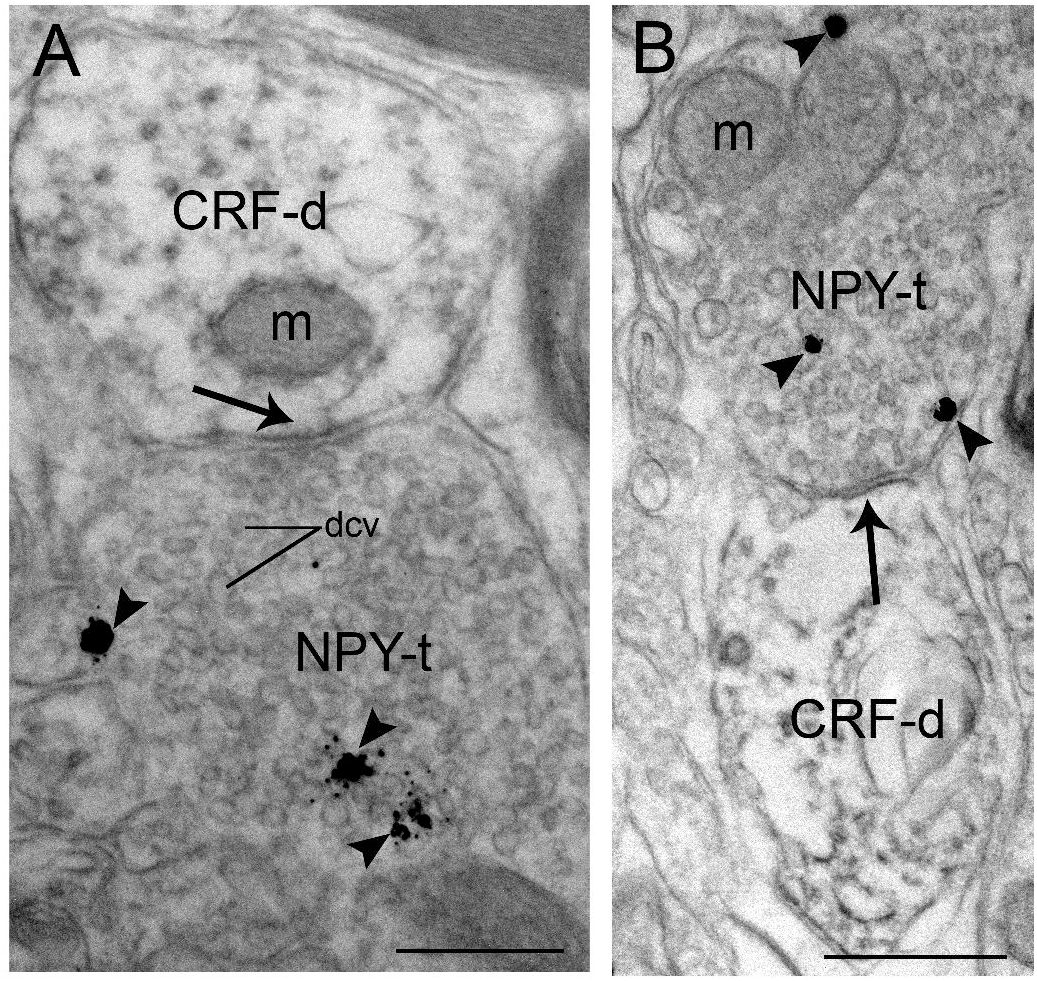
Electron photomicrographs showing reverse immunolabeling, in which neuropeptide Y (NPY) is labeled with immunogold-silver while corticotropin-releasing factor (CRF) is labeled with immunoperoxidase in the central nucleus of the amygdala (CeA). **A-B.** In each panel, an NPY-labeled axon terminal (NPY-t) forms symmetric type synaptic contact (arrow) with a CRF-labeled dendrite (CRF-d). The NPY-labeled axon terminal contains numerous small clear vesicles and dense core vesicles (dcv). Scale bar = 0.4 µm. Arrowheads point to immunogold-silver particles indicative of NPY immunoreactivity. m, mitochondria.

## Discussion

The current study presents, to our knowledge, the first anatomical evidence through both immunofluorescence and high-resolution anatomical analyses that NPY fibers establish direct contacts with neurons possessing the molecular components required for CRF release. These anatomical findings provide a cellular basis for interactions between the anti-stress neuropeptide NPY and the stress-associated neuropeptide CRF. This interaction may contribute to the response of the central amygdalar system to various stressors, and play a role in modulating the diverse functions attributed to the CeA. The present results suggest that NPY can directly influence or regulate CRF activity in the CeA, potentially serving as a modulatory mechanism through which NPY dampens or fine tunes the stress response mediated by CRF, an effect that may be pivotal in understanding stress-related psychiatric disorders and substance use disorders, and valuable for developing targeted treatments. The current study also established that individual NPY axon terminals co-express CRF, in addition to individual NPY axon terminals contacting CRF dendrites. This provides subcellular evidence that the regulation of the CRF-induced stress signaling pathway may be more complex than previously understood, and likely involves the simultaneous release of NPY as a coregulator during CeA regulation.

### Methodological considerations

The investigation of synaptic associations using immunohistochemical labeling combined with electron microscopy offers several advantages. For example, the pre-embedding immunohistochemical technique preserves the fine morphological details necessary to localize the antigen of interest at the subcellular level. The dual immunolabeling method, combining immunoperoxidase with immunogold-silver labeling, enables the identification of synaptic specializations within defined neuronal populations. However, pre-embedding dual immunolabeling approaches have limitations, particularly regarding the penetration of primary and secondary antibodies in thick tissue sections. Inadequate antibody penetration for CRF and NPY may lead to an underestimation of their distribution frequencies and the number of synaptic contacts on CRF-labeled dendrites. This limitation was carefully mitigated through steps previously described in the Methods section.

The quantification of dual-labeled neurons in the CeA is limited in the present study, as the analysis was conducted on tissue sections from non-colchicine-treated mice. Consistent with our previous reports (Reyes et al., 2015; Reyes et al., 2019), CRF-immunoreactive neurons were abundantly distributed in the CeA, as revealed by immunofluorescence. We acknowledge that colchicine pretreatment enhances CRF immunoreactivity in neuronal somata by disrupting microtubule-based peptide transport to axons, thereby causing an accumulation of labeling in the soma. However, previous studies have indicated that the presence or absence of colchicine can influence the apparent number of CRF neurons, potentially leading to underestimation (Van Bockstaele *et al*., 1998). Despite this, CRF neurons were clearly visible at both the immunofluorescence and ultrastructural levels across all profiles, suggesting that the absence of colchicine should not significantly impact the interpretation of our findings. Additionally, immunofluorescence demonstrated that NPY immunoreactivity was distinctly distributed in the processes and fibers of the CeA. Notably, the absence of colchicine facilitated better localization and visualization of NPY, particularly in axon terminals. While identifying CRF localization in dendrites is essential, our study also aimed to examine the localization of NPY in axon terminals to elucidate CRF-NPY interactions. Therefore, the dual-labeled neurons identified in this study should be considered an approximation and may underestimate the actual number. Importantly, the primary objective of this study was to define CRF-labeled amygdalar neurons that are contacted by NPY-labeled axons, rather than to quantify neuropeptide expression levels. Furthermore, it is worth noting that the use of colchicine could potentially alter the localization of NPY in axon terminals, further justifying our methodological choice.

### NPY modulation of CRF CeA neurons

The CeA is recognized as the primary output nucleus of the amygdala and is composed of highly intricate microcircuitry that mediates a wide range of responses to emotion, arousal and stress (Adhikari, 2014; Paretkar & Dimitrov, 2018). Consistent with previous reports in mice, CeA presents numerous CRF containing neurons (Asan et al., 2005; Wang *et al*., 2011) which mediate activation of CeA in response to diverse types of stressors (Mamalaki *et al*., 1992; Iwasaki-Sekino *et al*., 2009; Ciccocioppo *et al*., 2014). CeA neurons express a high concentration of CRF and NPY that changes depending on conditions. For instance, following restraint stress, footshock or psychological stress, CRF mRNA levels are elevated in the CeA (Mamalaki *et al*., 1992) while higher NPY levels are correlated with lower anxiety-like behavior (Sharko *et al*., 2016). Previous studies suggest that these neuropeptides play a key role in the amygdala’s influence on emotional responses (Heilig, 2004; Bowers *et al*., 2012).

Our present study showed at the ultrastructural level that NPY axon terminals contact CRF neurons. It is possible that these CRF neurons targeted by NPY axon terminals project to other brain regions involved in stress or substance use, as anxiety is a hallmark symptom of both stress-related psychiatric disorders (Al Bazzal et al., 2025) and substance use disorders (Walker, 2021). Efferent CRF neurons from the CeA are known to project to multiple brain regions including hypothalamus, brainstem and other amygdalar nuclei (Van Bockstaele et al., 1998; Van Bockstaele et al., 1999; Iwasaki-Sekino et al., 2009; Reyes et al., 2011; Beckerman et al., 2012; Reyes et al., 2017; Paretkar & Dimitrov, 2018; Reyes et al., 2019, Weera et al., 2023). Findings from the present study, supported by immunofluorescence and ultrastructural analysis, indicate that some of these CRF neurons in the CeA that project to these nuclei are contacted by NPY fibers. However, further studies are required to provide unequivocal evidence for this neuroanatomical pathway at the ultrastructural level. It would be particularly interesting to determine whether the population of CRF neurons in the CeA projecting to these nuclei is modulated by NPY during stress or following drug exposure. NPY-containing afferents to the CeA emanate from various brain nuclei including nucleus of the solitary tract, ventrolateral medulla and paraventricular thalamic nucleus (Zardetto-Smith & Gray, 1990; Freedman & Cassell, 1994; Zardetto-Smith & Gray, 1995). These findings provide a neuroanatomical pathway that underlies the cellular interactions between NPY and CeA neurons. Further research in needed to identify the sources of the NPY afferents to the CeA observed in the present study.

NPY regulates CRF neurons in the CeA primarily post-synaptically. Our current results demonstrate that the majority of NPY-containing axon terminals form symmetric synapses with CRF-containing dendrites indicating that NPY regulates CRF neurons via inhibitory neurotransmission. Notably, nearly all synapses formed were inhibitory and only very few were excitatory. The finding seems to contradict previous reports indicating that NPY axon terminals form asymmetric synapses with CRF-containing dendrites and soma in the paraventricular nucleus of the hypothalamus (PVN) (Liposits *et al*., 1988), demonstrating stimulatory effects of NPY on neuroendocrine responses. The differential effects of NPY on CRF neurotransmission in amygdalar and hypothalamic circuit may arise from differential coupling to effector systems or the differential distribution of NPY receptors on amygdalar or hypothalamic CRF neurons (Heilig *et al*., 1993; Dimitrov *et al*., 2007; Tasan *et al*., 2010; Pleil *et al*., 2015). It is further possible that the mechanism of how NPY modulates the CRF neurons activity in the CeA involves a circuit distinct from that used in the PVN. This is particularly noteworthy given the roles of both the CeA and PVN in stress-related neural circuitry. Further studies are required to clarify the mechanisms underlying the differential effects of NPY on CRF neurotransmission in the amygdala and hypothalamus.

Although CRF plays a central role in orchestrating the stress response, NPY acts to counterbalance the action of CRF (Sah & Geracioti, 2013; Sah *et al*., 2014; Sabban *et al*., 2015). The modulatory effects of NPY in the CeA are mediated through its interaction with NPY receptors. While NPY receptor activation leads to a reduction in cAMP production, CRF activation promotes cAMP production. Sheriff and colleague (Sheriff *et al*., 2001) employed an engineered immortalized amygdalar cell line (AR-cell), which expresses CRF (2alpha), Y1 and Y5 receptors as verified by RT-PCR to examine these opposing effects. They reported that treating AR-5 cells with 3 μM CRF resulted in a significant, concentration-dependent increases in cAMP production (Sheriff *et al*., 2001). In contrast, increasing concentrations of NPY markedly suppressed this CRF-induced cAMP production compared to the CRF treatment alone. These findings suggest that NPY may antagonize CRF’s effects by modulating the protein kinase A signaling pathway (Sheriff *et al*., 2001). Y1 receptor is one of the well characterized NPY receptor subtypes in the central nervous system and is predominantly found at postsynaptic sites, where it is thought to modulate neuronal activity (Kopp et al., 2002). In addition to Y1, Y2 receptor is a potential key player in the interaction between NPY and CRF in the CeA. In contrast to Y1 receptors, which are mainly postsynaptic, Y2 receptors are localized presynaptically acting as autoreceptors, and functionally opposing Y1 receptor activity (Parker & Balasubramaniam, 2008). Once Y2 receptors on NPY-containing terminals are activated, endogenous NPY release is suppressed (King et al, 1999), thereby reducing the availability of NPY in the synaptic cleft. Using elevated plus maze (EPM), the anxiogenic effects of Y2 receptors have been demonstrated (Nakajima et al., 1998). However, antagonism the Y2 receptors during abstinence completely reverses nicotine-induced social anxiety-like behavior, as shown in the social interaction test (Aydin et al., 2011) and alleviates the anxiogenic effects of alcohol withdrawal, as demonstrated in the EPM (Cippitelli et al., 2010; Rimondini et al., 2005). Among various amygdalar nuclei, the highest expression levels of Y2 receptors have been observed in the CeA (Wood *et al*., 2016).

In the CeA, NPY immunoreactivity is primarily observed in axon terminals, suggesting that presynaptic release of NPY may modulate stress-associated CRF activation of CeA neurons. Our present results show evidence that NPY and CRF are co-localized in the same presynaptic profile. This presents an intriguing cellular basis for interaction, as the functional impact for NPY released from axon terminals containing only NPY may differ from the NPY release from axon terminals containing both NPY and CRF. Further anatomical studies are required to elucidate the nature and significance of this of this interaction.

### Functional implications

The CeA is a key structure involved in emotional processing, arousal and stress regulation (Adhikari, 2014; Paretkar & Dimitrov, 2018). The current findings indicate an interaction between limbic CRF and NPY circuits, which may play an important role in the modulation of anxiety, fear, the stress response and autonomic regulation functions that are closely associated with CeA. As the primary output region of the amygdala, the CeA connects extensively with endocrine and autonomic centers in the hypothalamus and brainstem, enabling it to mediate both behavioral and physiological responses to emotional stimuli (Tsukioka *et al*., 2022). In other words, the current results also suggest that the NPY and CRF interactions within the CeA are essential in modulating emotional responses and autonomic regulation. For example, reduced NPY levels in the amygdala, along with elevated CRF levels in the CeA, are linked to the development of social anxiety behavior following nicotine exposure and subsequent abstinence (Aydin et al., 2011). In fact, treatment with an Y2 receptor antagonist reverses the nicotine-induced reduction in NPY levels and decreases CRF mRNA expression in the CeA of nicotine-pretrained cohorts exhibiting high locomotor response to novelty, restoring both to levels comparable to those of saline-pretrained controls (Aydin et al., 2011). These findings suggest that the interaction between NPY and CRF in the CeA plays a critical role in modulating negative affect, such as anxiety, particularly during drug withdrawal. It can be speculated that the interaction between NPY and CRF may modulate gamma-aminobutyric acid (GABA) neurotransmission. For example, whole-cell recordings in slices of the bed nucleus of the stria terminalis have shown that NPY inhibits, while CRF enhances, GABAergic neurotransmission. In this nucleus, pharmacological studies have shown that NPY attenuates GABAergic neurotransmission via activation of Y2 receptors, whereas both pharmacological and genetic evidence suggest that CRF and urocortin enhance GABAergic neurotransmission via activation of the CRF receptor type 1 (Kash & Winder, 2006).

Additionally, dysregulation of NPY signaling within the CeA is increasingly recognized as a contributing factor to the pathophysiology of various stress-related disorders (Eaton *et al*., 2007; Sah & Geracioti, 2013; Sah *et al*., 2014; Sabban *et al*., 2015; Serova *et al*., 2019), further underscoring the relevance of such interaction as a promising target for therapeutic intervention. The NPY system is activated following stress exposure and contributes to resilience by mitigating the harmful effectss of stress (Sah & Geracioti, 2013; Sah *et al*., 2014; Sabban *et al*., 2015; Sabban *et al*., 2016). Pharmacological research in rodents highlights the crucial role of NPY in promoting anxiolysis and dampening anxiety-like behaviors. NPY administration intracerebroventricularly in rodents has been shown to produce anxiolytic effects across multiple paradigms of anxiety-like behavior including elevated plus-maze, open field test and fear-potentiated startle test (Broqua *et al*., 1995; Sorensen *et al*., 2004). Similarly, direct administration of NPY into the amygdala reduces anxiety-like behaviors and promotes anti-depressant effects (Sajdyk *et al*., 2002; Taksande *et al*., 2014). Genetic research has validated the anxiolytic role of NPY in rodents. Mice lacking the NPY gene exhibit increased anxiety-like behavior (Bannon *et al*., 2000) whereas overexpression of NPY in the amygdala via a recombinant adeno-associated viral vector results in marked anxiolytic-like effects across various anxiety-related behavioral tests (Christiansen *et al*., 2014). Studies in rodents show that intranasal delivery of NPY administered either prior to or immediately after exposure to traumatic stress prevented the development of behavioral, neuroendocrine, and molecular dysfunctions characteristic of PTSD as these studies used the single prolonged stress (SPS) model of PTSD (Serova *et al*., 2014; Sabban *et al*., 2015; Sabban *et al*., 2016). Studies in humans show that lower NPY levels are correlated with anxiogenic trait, increased emotionality, and reduced resiliency to stress (Zhou *et al*., 2008). For example, among soldiers undergoing military training, lower NPY were associated with increased distress and poorer stress-related performance while higher NPY levels were linked to greater resilience and improved performance under stress (Morgan *et al*., 2000). Furthermore, higher NPY levels in veteran soldiers were associated with better coping and recovery from the adverse effects of stress (Yehuda *et al*., 2006). Our current findings that CRF neurons in CeA are innervated by NPY at both immunofluorescence and ultrastructural levels unravel the position of NPY in modulating CRF signaling in the CeA.

## Acknowledgements

This research was supported by the National Institute on Alcohol Abuse and Alcoholism of the National Institutes of Health (NIH) under Award Number R21AA030361 (to BASR and JMB). The content is solely the responsibility of the authors and does not necessarily represent the official views of the NIH.

